# Genome-wide epistasis and co-selection study using mutual information

**DOI:** 10.1101/523407

**Authors:** Johan Pensar, Santeri Puranen, Neil MacAlasdair, Juri Kuronen, Gerry Tonkin-Hill, Maiju Pesonen, Brian Arnold, Yingying Xu, Aleksi Sipola, Leonor Sánchez-Busó, John A Lees, Claire Chewapreecha, Stephen D Bentley, Simon R Harris, Julian Parkhill, Nicholas J Croucher, Jukka Corander

## Abstract

Discovery of polymorphisms under co-selective pressure or epistasis has received considerable recent attention in population genomics. Both statistical modeling of the population level co-variation of alleles across the chromosome and model-free testing of dependencies between pairs of polymorphisms have been shown to successfully uncover patterns of selection in bacterial populations. Here we introduce a model-free method, SpydrPick, whose computational efficiency enables analysis at the scale of pan-genomes of many bacteria. SpydrPick incorporates an efficient correction for population structure, which is demonstrated to maintain a very low rate of false positive findings among those SNP pairs highlighted to deviate significantly from the null hypothesis of neutral co-evolution in simulated data. We also introduce a new type of visualization of the results similar to the Manhattan plots used in genome-wide association studies, which enables rapid exploration of the identified signals of co-evolution. Application of the method to large population genomic data sets of two major human pathogens, *Streptococcus pneumoniae* and *Neisseria meningitidis*, revealed both previously identified and novel putative targets of co-selection related to virulence and antibiotic resistance, highlighting the potential of this approach to drive molecular discoveries, even in the absence of phenotypic data.

## INTRODUCTION

Statistical analysis of co-variation between non-adjacent sites in large protein alignments has matured since its inception, over 20 years ago (*1–7*). More recently, attention has also been directed towards performing a similar type of exploratory analysis of genome-wide nucleotide alignments for bacterial populations to reveal putative sites evolving under co-selective pressures and possibly being involved in epistatic interactions (*8–10*). Genome-scale analysis of co-variation at single-nucleotide resolution, here termed as genome-wide epistasis and co-selection study (GWES), poses considerable computational challenges as the number of pairs to be considered increases quadratically with the number of sites. Previous GWES approaches have been based on either straightforward pairwise tests (*8*), which do not distinguish between indirect and direct interactions, or a more elaborate model-based technique known as direct coupling analysis (DCA) (*9, 10*).

The main motivation behind pairwise methods has typically been scalability, however, a recent simulation study on high-dimensional structure learning of synthetic network models showed that a family of pairwise methods based on mutual information (MI) may be as accurate as and even outperform model-based methods in the small sample regime (arXiv:1901.04345), which is the typical setting for most bacterial population genomic data. While MI has been proposed for the analysis of protein alignments (*1, 11*), it has not yet been systematically applied to bacterial population genomics. Here we introduce a novel MI-based GWES method, SpydrPick, which is scalable to an order of magnitude larger data sets than those considered so far in DCA-based GWES (*9, 10*).

To account for population structure, we use a sequence reweighting technique commonly employed when analysing protein sequence alignments (*3, 4, 11*), and also more recently when performing GWES (*9, 10*). However, a different route is taken towards selecting the best candidates of directly co-selected or interacting mutations among the identified signals of co-variation. These are chosen as the significant outliers in terms of a global background distribution estimated across the genome, combined with a pruning method introduced for analyses of gene expression data (*12*). The focus on the statistical quantification of the background pattern across the genome lends itself well to an intuitive and efficient visualization of the results akin to a Manhattan plot used in genome-wide association studies, which we term as the GWES Manhattan plot.

We demonstrate the usefulness and reliability of SpydrPick by application to both simulated sequences evolving under a neutral model, and to two large population genomic data sets of the major human pathogens *Streptococcus pneumoniae* and *Neisseria meningitidis*. For the latter pathogen, we analysed the entire pan-genome, which contains so many mutations that most model-based approaches are computationally infeasible, including even the recent highly optimized DCA-based software (*10*).

## MATERIAL AND METHODS

### Method

An overview of the SpydrPick pipeline is shown in Figure 1. The different steps are described in detail in the following sections.

**Figure 1.**
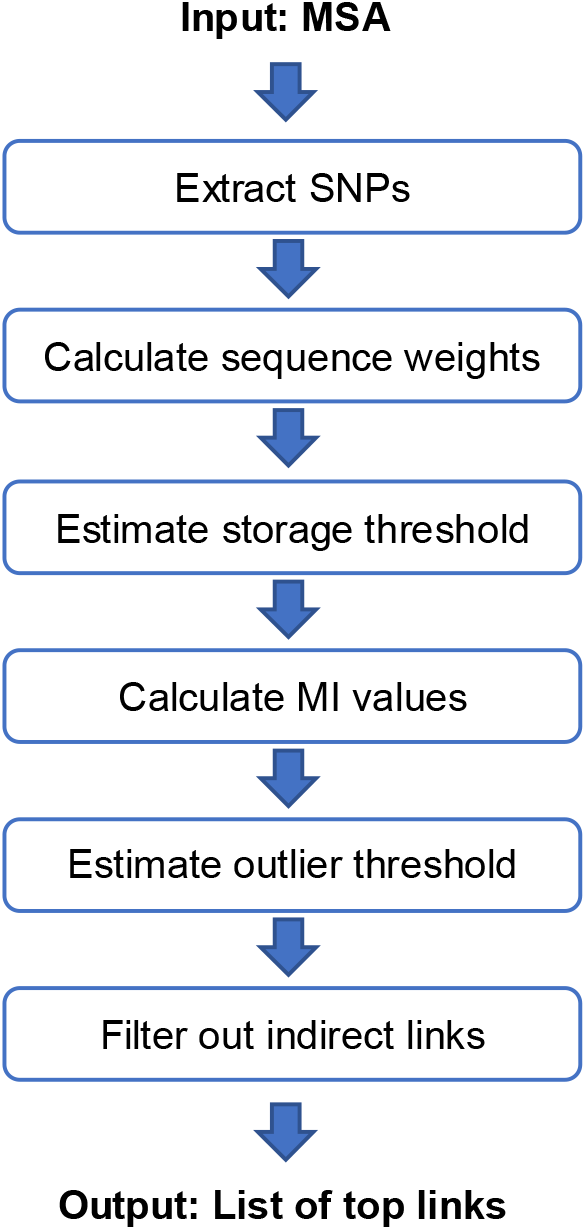
An overview of the SpydrPick pipeline.

*Mutual information*. Mutual information (MI) is an information theoretic measure of the mutual dependence between two random variables. More specifically, let *X* and *Y* be two discrete random variables with outcome spaces *val*(*X*) and *val*(*Y*). The MI between *X* and *Y* is then formally defined by

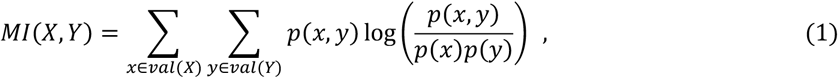

where *p*(*x,y*) is the joint probability of *X* = *x* and *Y* = *y*, while *p*(*x*) = Σ_*y*∈*val*(*Y*)_*p*(*x,y*) and *p*(*y*) = Σ_*x*∈*val*(*X*)_*p*(*x,y*) are the corresponding marginal probabilities. In practice, the joint distribution is typically unknown and has to be estimated from data. Let *n*(*x,y*) denote the count of the joint outcome *X* = *x* and *Y* = *y* occurring in a data set containing *n* independent and identically distributed (IID) observations generated from *p*(*X,Y*). Typically, the joint probabilities are estimated by the relative frequencies of the joint outcomes corresponding to maximum likelihood estimates. To avoid issues related to zero counts and increase the stability of the estimator, we add 0.5 to the joint counts according to

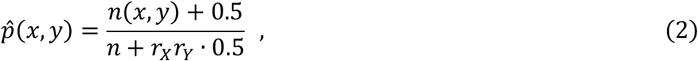

where *r_X_* and *r_Y_* denote the number of possible outcomes for *X* and *Y*, respectively. In the Bayesian framework, the above point estimator corresponds to the posterior mean under a Dirichlet prior distribution with the hyperparameters set to 0.5, corresponding to Jeffreys’ prior (*13*).

#### Sequence reweighting

In the context of this work, *X* and *Y* in the previous paragraph correspond to single-nucleotide polymorphisms (SNPs) and the outcome spaces represent the four nucleotides *A,C,G,T* with an additional outcome representing gaps. The observed data is in form of a multiple sequence alignment (MSA) containing *n* sequences (*S*_1_,…,*S_n_*) of length *L*. In general, the sequences in an MSA strongly violate the IID assumption since they share a linkage through an evolutionary relationship. This is problematic from a practical point of view, since potentially interesting signals may be hidden behind background noise caused by the population structure within an MSA. Consequently, to adjust for the population structure in the MI estimator, we apply a technique known as sequence reweighting, which has successfully been used previously for both protein contact prediction (*3, 4*) and GWES (*9, 10*). Reweighting assigns a weight to each sequence according to how different it is from the other sequences in the MSA, such that the counts of allele pairs occurring in the MI estimator will reflect the level of clusteredness across the MSA.

Let *m_i_* denote the number of sequences (including *S_i_*) whose mean per-site Hamming distance to *S_i_* is smaller than a similarity threshold, for which we used a default value of 0.10. The weight given to sequence *S_i_* is then calculated by

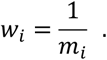

The effective count *n*_eff_(*x,y*) is calculated by summing the weights of all sequences with the corresponding joint configuration over the SNP sites represented by *X* and *Y*. The counts in (*2*) are then replaced with the corresponding effective counts:

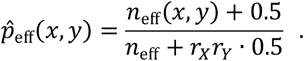

The above estimates are finally plugged into (*1*) resulting in the reweighted MI estimator.

#### Filtering out indirect links

An unavoidable issue with methods based solely on pairwise association tests is their inability to distinguish between direct and indirect associations. In particular, in the GWES context it is typically expected that a strong direct dependence between two distant SNP sites would be accompanied by a collection of slightly weaker indirect dependencies between sites in close proximity of the coupled sites due to genetic linkage. As a result, pinpointing the exact locations of co-evolving loci at SNP resolution in a bacterial GWES is in general very difficult due to strong linkage disequilibrium between nearby sites. Still, considering that the identified links need to be examined manually, our aim is to produce as compact a list of SNP pairs as possible, containing the most likely candidates of mutations co-evolving under a shared selection pressure.

To select a subset of SNP pairs containing only the most promising links, we use the same filtering technique as ARACNE, which was originally introduced as a method for inferring gene expression networks (*12*). The filtering technique is based on a property known as the data processing inequality, which states that if two variables *X* and *Y* only interact through a third variable *Z*, then

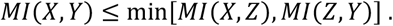

In other words, the indirect dependence between *X* and *Y* cannot be larger than either of the two direct dependencies through which it is mediated. Formally, ARACNE starts from a graph containing a link for each non-zero MI value. The algorithm then examines each triplet of mutually linked variables and removes the weakest link (see Figure 2). In the degenerate case, where there is no unique weakest link in a triplet, no link is removed. The algorithm is order-independent in the sense that a link that has been marked for removal from one triplet is still considered present with respect to a non-examined triplet containing that link.

**Figure 2.**
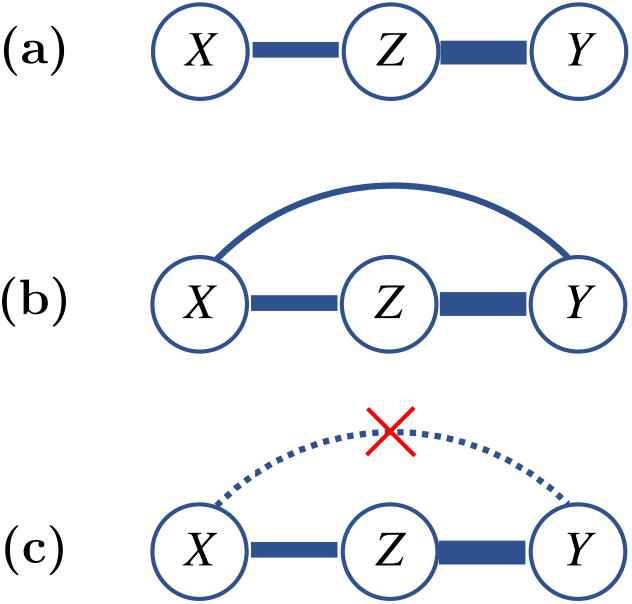
Illustration of the ARACNE step (the width of the links represents the interaction strength): (a) True interaction structure: *Z* is strongly linked to *X* and Y, which are not directly linked to each other. (b) A pairwise test outputs a significant association between *X* and *Y* due to the indirect link through *Z*. (c) The ARACNE step removes the indirect link between *X* and *Y*, being the weakest out of the three links.

Naively applying the ARACNE filtering step would be computationally intractable, since there are in total 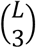 possible triplets. However, in practice it is sufficient to run the procedure over a small list containing only the top estimated links. Consequently, the main computational part will still be to estimate the MI values over the 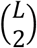 pairs. The ARACNE approach is not only appealing due to its computational simplicity, but also its ability to produce a small representative set of links that are most likely to be direct. One of the drawbacks with this approach is that it will never output a triplet of mutually linked sites (except in the degenerate case) even if such a triplet existed. However, three mutually linked sites will still be contained in a single connected component and thus the association between the three loci will remain visible.

#### Threshold for result storage

Saving the complete output of a GWES to disk would typically result in such large files that they would become unwieldy. Nevertheless, since the main target is to identify the largest MI values, estimation results can be filtered online (i.e. as each new value is calculated) to reduce the amount of storage required. To this end, we use a subsampling procedure to determine a threshold for saving a user-specified top fraction of the MI values. This is done by randomly selecting a subset of SNP pairs for which the MI values are calculated. The empirical cumulative distribution function is then used to estimate an appropriate saving threshold that corresponds to the user-specified top fraction. To increase stability, the procedure is repeated several times and the median threshold value is selected for final filtering.

#### Outlier analysis

To assess if a link is strong enough to warrant further study, we perform an outlier analysis. Due to genetic linkage, SNPs in close chromosomal proximity tend to be in strong linkage disequilibrium (LD). Note that LD here refers to SNPs showing a significant association specifically due to close genetic linkage. Since strong LD masks any potential signal of shared co-evolutionary selection pressure, we restrict the outlier analysis to non-LD pairs. The default approach for filtering out LD-pairs is to use a simple distance-based cut-off. For this, we used a default cut-off value of 10 kbp.

To estimate an outlier threshold among the non-LD pairs, we use a data-driven procedure based on Tukey’s outlier test (*14*). The test assesses how extreme an MI value is in comparison to a global background distribution observed for the analysed data set. If the MI value of a direct link is flagged as an outlier, the corresponding SNP pair will automatically be carried forward for further analysis. As background distribution for the outlier test, we use an extreme value distribution by which we effectively attempt to model the distribution of maximum MI values for a site (w.r.t. non-LD pairs). In practice, we save the maximum MI value of each site and calculate the lower (*Q*_1_) and upper (*Q*_3_) quartiles of empirical extreme value distribution. Following Tukey’s criterion, we then label an MI value larger than *Q*_3_ + 1.5 × (*Q*_3_ − *Q*_1_) as an outlier. In addition to the default threshold, we label an MI value larger than *Q*_3_ + 3 × (*Q*_3_ − *Q*_1_) as an extreme outlier.

The typical approach for determining significance in this type of problem is to run a permutation analysis (*12, 15*). For this application, such an approach would be too inclusive since the maximum MI values observed in the background distribution of real MSAs exceed those observed under a null model in which the sites are unlinked through permutations. Moreover, the extent of the tail region of the background distribution may vary significantly between data sets due to differences in population structure, recombination rate, etc. For this reason, our significance analysis is based on identification of outliers among the actual MI values observed for a particular population. Being based on quartiles, Tukey’s outlier test is by design very robust against extreme values. The critical assumption behind this procedure is that the majority of SNPs are not linked to other SNPs beyond LD.

#### MutuaI information without gaps

When calculating the MI values, gaps are by default considered an outcome. While some gaps can be informative, others may simply be due to difficulties in the sequencing process: difficult-to-sequence regions may be systematically absent from all lower-quality sequences, resulting in distinct patches of gap characters that appear in parallel across samples. Hence, some interactions may be artificially amplified in regions with low-quality sequence data. To facilitate discovery of such cases in the subsequent manual analysis, we also calculate the MI value of the top pairs using only sequences where neither site of a pair contains a gap. Since the collection of sequences without gaps varies between pairs, it is difficult to compare gap-free MI values between SNP pairs in a meaningful way, however, the gap-free MI value can still be informative for a given pair in the sense that a large decrease in MI when dropping the gap sequences is an indication of a gap-driven interaction.

#### Implementation

SpydrPick was implemented in C++ and supports parallel execution in a shared memory environment. Its space-efficient data structure, indexing strategy and online filtering of output jointly enable excellent scalability to an order of magnitude larger genome data sets than previous software developed for epistasis and co-selection analysis.

#### GWES Manhattan plot

For compactly visualizing the results of a GWES, we use a modified version of the GWAS Manhattan scatter plot. In a standard GWAS Manhattan plot, the association strength between a SNP and some phenotype (y-axis) is plotted against the chromosomal location of the SNP (x-axis), meaning that each point represents a single SNP. A GWES Manhattan plot has a similar design, however, each point now represents a pair of SNPs such that the x-axis displays the distance between the chromosomal locations of the SNPs and the y-axis displays the association strength between the SNPs, which is determined by their MI value.

### Data

#### Neutral model

To ensure that the designed method maintains a sufficiently low rate of false positive findings indicated as outliers, we generated genomic data with realistic LD under a neutral population model using the population simulator introduced in (*16*). Thus, the simulation illustrates the output of the method in a controlled, yet challenging scenario where there are no co-evolving SNP pairs beyond the LD pattern imposed by the neutral model.

The genome was a linear chromosome of 200 kbp and the parameters of the simulator model were set to represent a challenging heavily structured population with 20 inter-connected subpopulations, (see Table S1 for exact simulator settings). The simulation was repeated ten times with different random seeds. From each population of 20,000 isolates, a random sample covering 5% of the population was drawn, resulting in 886 – 912 unique sequences per sample. The simulated alignments were filtered for bi- and multi-allelic loci with a minor allele frequency (MAF) greater than 1% and a gap frequency (GF) smaller than 15%. The number of SNPs per filtered alignment was in the range of 10,568 – 12,400. For one of the simulated alignments, a phylogenetic tree was estimated using RAxML with the default settings and GTR+Gamma model (*17*).

#### Streptococcus pneumoniae

Our first real alignment contained 3,042 *S. pneumoniae* strains collected in Maela, a refugee camp close to the border between Thailand and Myanmar (*18*). The whole genome alignment was generated from short-read data aligned to the reference sequence of *S. pneumoniae* ATCC 700669 whose genome is a circular chromosome of 2,221,315 bp (*19*). For the GWES, bi- and multi-allelic loci with MAF greater than 1% and GF smaller than 15% were included in the analyses. The filtered alignment contained 94,880 SNPs.

The diverse population structure in the data, together with the recombinant nature of *S. pneumoniae*, make the data ideal for GWES (*9*). Moreover, this particular data set has previously been analysed by DCA approaches, which successfully discovered several interacting regions with plausible biological explanations (*9, 10*). Hence, the main aim for this data set was to investigate how well the earlier highlight findings could be rediscovered using our model-free method.

#### Neisseria meningitidis

Our second real alignment contained 2,148 *N. meningitidis* strains, of which 543 were published by Lucidarme *et al*. 2015 (*20*) and the rest were obtained from different sequencing projects run in the Wellcome Sanger Institute, Cambridge (see Table S4 for more details). The pan-genome of the strains included in the study was created using Roary (*21*), with a percentage of isolates needed to consider a gene as core set to 95%. The core gene alignment and individual gene alignments of the 13,052 genes conforming the pan-genome under the above criteria were obtained directly from the output. All individual genes were concatenated to obtain a pan-genome-wide alignment of 11,375,926 bp using the Alignment Manipulation and Summary (AMAS) tool (*22*). For the GWES, bi- and multi-allelic loci with MAF greater than 1% and GF smaller than 70% were included. The filtered alignment contained 137,814 SNPs. An approximately-maximum likelihood phylogenetic tree was estimated with FastTree (*23*) from the SNP sites in the core alignment (obtained with SNP-sites (*24*)) using the GTR model of nucleotide substitution and gamma rate heterogeneity among sites.

In contrast to the *S. pneumoniae* alignment, where all sequences were mapped to a reference sequence, this pan-genome-wide alignment was constructed by concatenating individual gene alignments. As a result, we can no longer use a straightforward distance-based cut-off to filter out LD-mediated links. Instead, we simply define two sites within the same gene as an LD-pair and two sites from different genes as a non-LD pair. The main aim for this data set was to investigate if our method would still be able to extract plausible signals of co-selection under this modified setup.

## RESULTS

### Neutral model

The complex structure of the population generated under the neutral model is visible in the estimated phylogenetic tree, which has a large number of well separated clades (Fig 3a). High clonality within the clades is reflected by a low effective sample size, *n*_eff_ = 16.22, which is only 1.8% of the original sample size, *n* = 897. The Manhattan plot illustrating the output of SpydrPick is shown in Fig 3b. For short-distance SNP pairs we observe a peak in MI values due to LD. As the distance increases, the background distribution flattens out and remains at a constant level. The LD threshold at 10 kbp is marked with a red vertical line. The lower and upper horizontal red lines in the plot mark the outlier and extreme outlier threshold, respectively.

**Figure 3.**
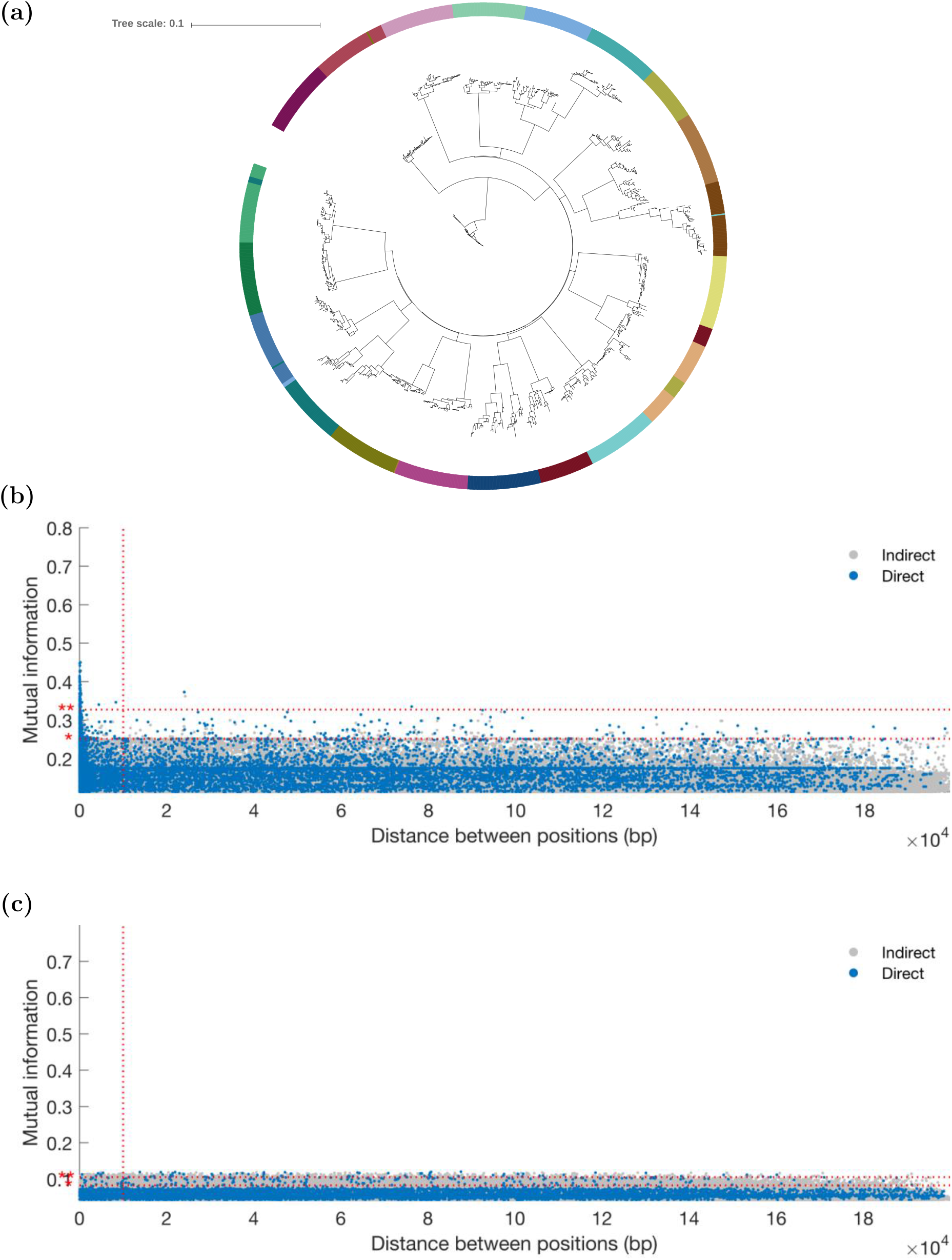
Neutral model - (a) Phylogenetic tree, (b) GWES Manhattan plot, (c) GWES Manhattan plot when the positions have been unlinked through permutations. (b) - (c) Direct and indirect links are plotted in blue and grey, respectively. The red horizontal dotted lines show the outlier thresholds; outlier * and extreme outlier **. The red vertical dotted line shows the LD threshold.

Blue points located right of the vertical line and above the horizontal line(s) can be considered false positives (FPs), since the simulator does not let any specific site patterns influence reproductive fitness. For the default outlier threshold, the average number of FPs over ten generated sequence alignments was 122.2 and the corresponding average FP rate was 2.1×10^−6^. For the extreme outlier threshold, the average number of FPs was 2.7 and the average FP rate 4.9×10^−8^. This shows that our method is able to maintain a low FP rate even under a very challenging population structure.

Finally, to illustrate the difference between the background distribution observed in Fig 3b and a corresponding null distribution, which was obtained by permuting the columns of the alignment used in Fig 3b, we have included a Manhattan plot of the SpydrPick output for the null distribution in Fig 3c. First, and as expected, the short-distance peak is no longer present in Fig 3c, since the permutation breaks the LD. Second, the level of the background distributions in Fig 3b clearly exceeds the corresponding null distribution in Fig 3c. As a result, any outlier threshold estimated from the permutation null distribution would likely be too inclusive with respect to the true background distribution, resulting in a high FP rate.

#### Streptococcus pneumoniae

After reweighting with respect to the filtered alignment, the effective sample size was reduced to *n*_eff_ = 130.26. The Manhattan plot of the analysis output is shown in Fig 4a. There is a high LD peak for short-distance pairs which eventually flattens out around 10 kbp (see Fig 4b) into a global background distribution. The striking difference from the simulated data is that there are now several distinct peaks clearly rising above the background distribution. Each peak is made up of a large collection of potential links. However, the ARACNE step filters out the vast majority as indirect, and only a few representative links (blue points) are singled out for further examination. In total, 163 direct links were flagged as outliers and 16 as extreme outliers. Here, we look closer at the extreme outliers, which are listed in Table S2. To facilitate the interpretation of the results, we have annotated the most interesting peaks in the Manhattan plot in Fig 4a using the distance column in Table S2. Finally, the Phandango plot (*25*) in Fig 5 shows the allele distributions across the population of the loci involved in the top links alongside phenotypic information about encapsulation and beta-lactam resistance.

**Figure 4.**
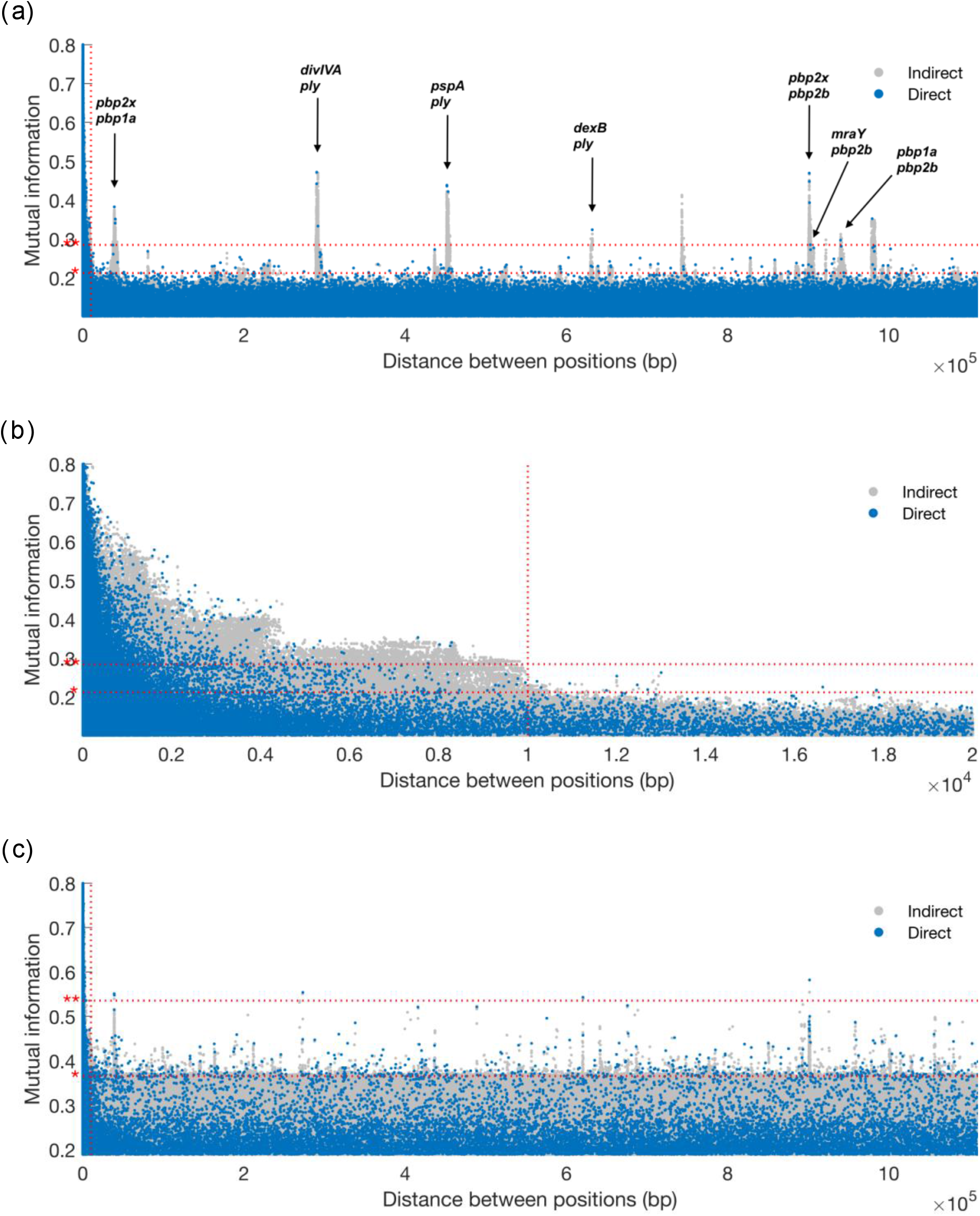
*S. pneumoniae* - GWES Manhattan plots: (a) complete distance range and with annotated peaks, (b) distances in the range 0 – 20 kbp, (c) complete distance range but without sequence reweighting. Direct and indirect links are plotted in blue and grey, respectively. The red horizontal dotted lines show the outlier thresholds; outlier * and extreme outlier **. The red vertical dotted line shows the LD threshold.

**Figure 5.**
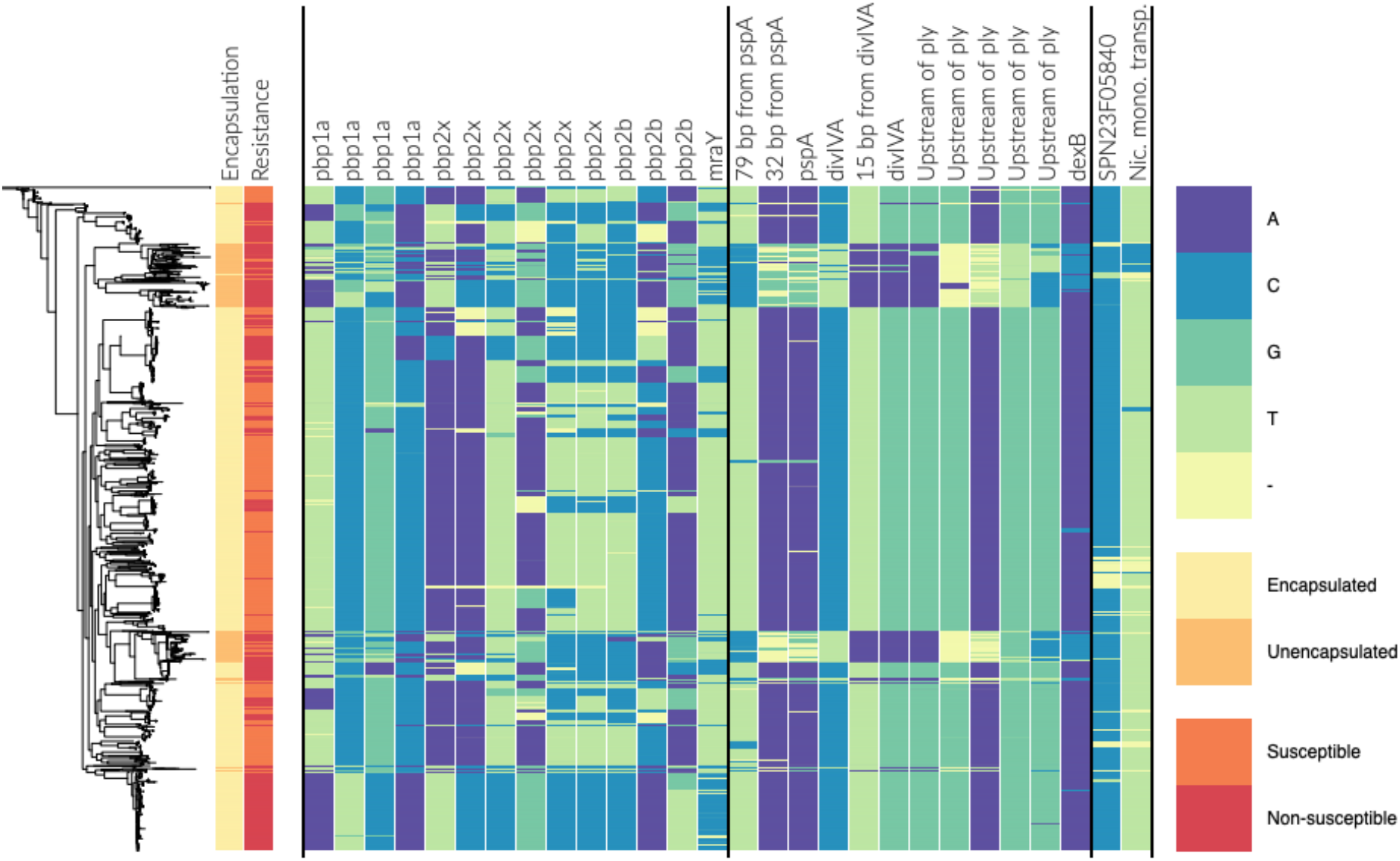
Phenotype information (encapsulation and beta-lactam resistance) and allele distribution at loci involved in the top links for the *S. pneumoniae* population. The estimated phylogeny is shown on the left. The two first columns are labelled by phenotype information and the remaining columns are labelled by gene name/id. The loci are sorted component-wise such that all columns within two successive vertical lines belong to the same component.

The majority of the top-ranking links discovered in the earlier DCA-based GWES (*9, 10*) were between three genes encoding penicillin-binding proteins (PBPs): SPN23F03410 (*pbp1a*), SPN23F16740 (*pbp2b*) and SPN23F03080 (*pbp2x*). These three proteins are involved in cell wall metabolism, and are the primary targets of beta lactam antibiotics. Modification of all three sequences is required for *S. pneumoniae* to exhibit high-level resistance to beta lactam antibiotics (*26–29*). Among the top 16 SpyderPick hits, 7 are between PBPs and the corresponding peaks are at distances 0.4×10^5^, 9.0×10^5^ and 9.4×10^5^ bp in the Manhattan plot. In addition to the links between the PBPs, there is also one link from *pbp2b* to SPN23F03090 (*mraY*), which is located directly downstream of *pbp2x*. The *mraY* gene encodes a phospho-N-acetylmuramoyl-pentapeptide-transferase also involved in cell wall biogenesis and, as noted by (*9*), it has been predicted that mutations in this transferase could be compensating for the costs of evolving beta lactam resistance (*29*).

In addition to the PBP-related links, there are 4 links involving SPN23F19490, which is part of the gene cluster SPN23F19480 - 19500 located directly upstream of SPN23F19470 (*ply*), encoding the toxin and key virulence factor pneumolysin. The ply-associated gene is coupled with SPN23F16620 (*divIVA*), SPN23F01290 (*pspA*) and SPN23F03150 (*dexB*), corresponding to the peaks at distances 2.9×10^5^, 4.3×10^5^ and 6.3×10^5^ bp in the Manhattan plot. The *divIVA* gene encodes a cell morphogenesis regulator and *pspA* encodes a surface protein associated with virulence. Links between ply-associated genes, *divIVA* and *pspA* were discovered as significant by the initial DCA method (*9*), but not by the more recent DCA method (*10*). A plausible reason for this is that the more recent DCA method fits a global model over all sites, whereas the initial DCA method uses a subsampling technique that makes it more similar to our local approach. The final link involving *dexB* has not been previously detected by any method. The *dexB* gene is located adjacent to the capsule polysaccharide synthesis locus in most *S. pneumoniae*, suggesting a possible link between the extracellular polysaccharide and the surface-associated PspA, Ply and DivIVA proteins. Further examination revealed the minor alleles at these loci were confined to several phylogenetically distinct clusters of non-typeable (unencapsulated) isolates, which lack a functional capsule polysaccharide synthesis locus (see Fig 5). This suggests non-typeable *S. pneumoniae* are not simply bacteria that have lost their capsule, but have also undergone other adaptive changes in specialising to a distinct niche. This may account for the distinct pathogenesis of unencapsulated strains, which do not cause severe invasive disease (*30*), but are known to cause outbreaks of conjunctivitis (*31*).

There are two remaining peaks in the Manhattan plot exceeding the extreme outlier threshold. The first peak at distance 7.4×10^5^ bp corresponds to an interaction between *pspA* and *divIVA*. This peak is not represented in the top links since its consistently the weakest link in triplets connecting *ply, pspA* and *divIVA*, and has therefore been labeled as indirect. The second and final peak is an example of a gap-driven signal. The MI of the corresponding link drops from 0.352 to 0.006 when excluding sequences that contain a gap on either site (see Table S2).

Finally, to illustrate the effect of the population structure, the result of running the analysis without sequence reweighting is shown in the Manhattan plot in Fig 4c. When comparing to the original plot in Fig 4a, it is clear that sequence reweighting is an essential step in separating the signal from the background distribution.

#### Neisseria meningitidis

After reweighting with respect to the filtered alignment, the effective sample size was reduced to *n*_eff_ = 515.86. The Manhattan plot of the analysis output for intra- and inter-gene pairs are shown in Figs 6a and 6b, respectively. Note that the distance between sites in Fig 6b is not a true distance, but a mock distance constructed for illustrative purposes from the ordering of the genes in the alignment. As expected, Fig 6a shows an abundance of high MI values among intra-gene pairs, especially among short-distance pairs. Fig 6b indicates that there is a collection of interaction signals rising above the global background distribution. Still, the overall signal-to-noise (or signal-to-background) ratio appears lower than in the *S. pneumoniae* analysis, which is also reflected by high outlier thresholds. A likely explanation for this is the inclusion of LD-mediated inter-gene links. In total, 48 direct links are flagged as outliers. In the following, we look closer at the 28 top-ranked links, which are listed in Table S3. The allele distributions of the loci involved in these links are visualized by the Phandango plot in Fig 7.

**Figure 6.**
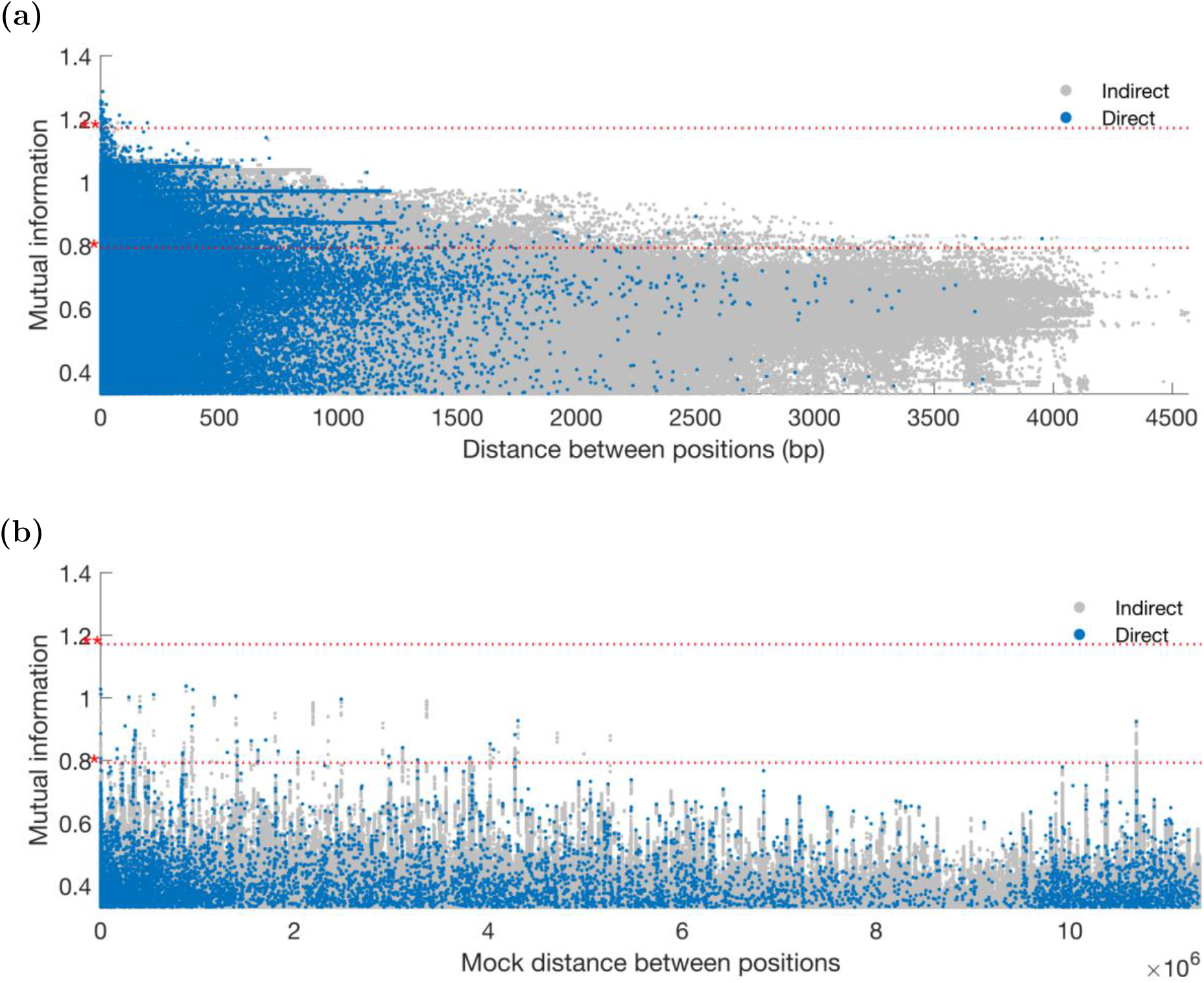
*N. meningitidis* - GWES Manhattan plots: (a) intra-gene links, (b) inter-gene links. The mock distance in (b) was calculated using the gene order in the actual alignment and is therefore not a true distance. Direct and indirect links are plotted in blue and grey, respectively. The red horizontal dotted lines show the outlier thresholds; outlier * and extreme outlier *.

**Figure 7.**
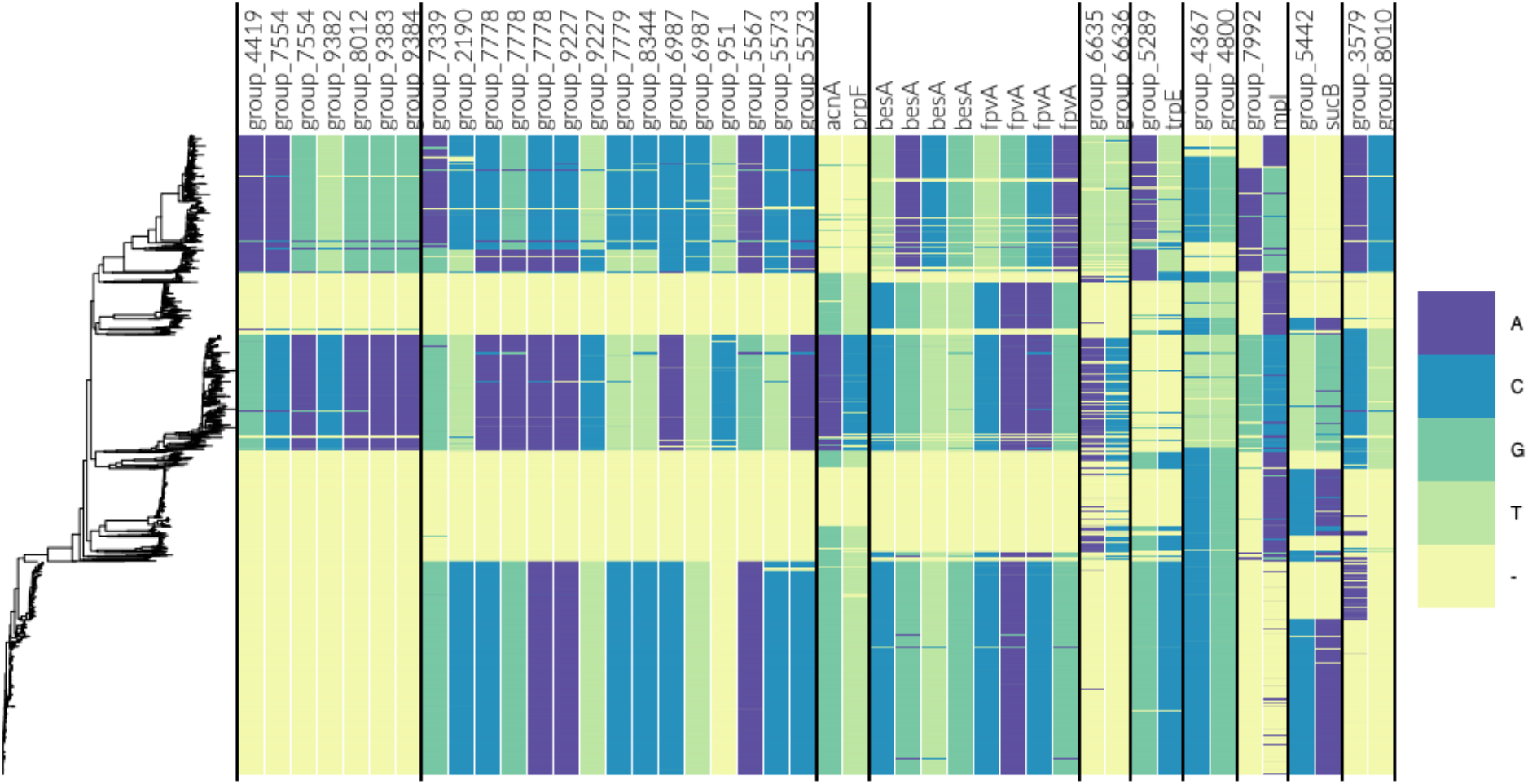
Allele distribution at loci involved in the top links for the *N. meningitidis* population. The estimated phylogeny is shown on the left and each column is labelled by gene name/id. The loci are sorted component-wise such that all columns within two successive vertical lines belong to the same component.

The majority of the identified links are between proteins of unknown function, many of which display high similarity to other phage-associated proteins or phage repressors. Previous work has identified a certain bacteriophage as important to virulence in *N. meningitidis* (*32, 33*), but the phage-associated proteins detected in this scan could not be further identified. To better assess the likelihood of LD causing the elevated MI values, we mapped the genes involved in the top links onto the reference genomes MC58 (*34*) and FAM18 (*35*), and calculated the inter-gene distances (Table S3). This revealed that most of the links were relatively short-distance, making it difficult to rule out the possibility of LD, especially for intra-phage-links. Hence, we looked closer at the 5 long-distance links for which the involved genes were more than 10 kbp apart in the reference genomes.

Out of the 5 long-distance links, 4 links were between the gene *besA*, encoding ferri-bacillibactin esterase, and the ferripyoverdine receptor *fpvA*. Both genes are involved in iron uptake during colonisation (*36, 37*). Iron uptake is an important pathway in most bacteria that colonise human hosts, and *Neisseria* is no exception, where iron uptake has been identified as an important determinant of virulence (*38, 39*), and essential for successful colonisation (*39*). The besA and fpvA genes are located 62,477 bp apart in the MC58 reference genome and 63,514 bp apart in the FAM18 reference genome, and the strong links are thus very unlikely to be caused by the background LD.

The final long-distance link is between the anthranilate synthase component I, *trpE*, involved in tryptophan synthesis, and a hypothetical gene, here referred to as *group_5289* (name given by Roary). When searched against the non-redundant protein database with tblastx (*40*), *group_5289* showed similarities with a betaine transporter. The *trpE* and *group_5289* genes are 361,849 bp apart in the MC58 reference genome and 722,196 bp apart in the FAM18 reference genome. From previous molecular biology work studying these pathways, we can see how these two genes might come to be under selection. Tryptophan synthesis is a crucial part of protein biosynthesis, and its synthesis has been linked to greater virulence in other bacterial species by allowing for immune evasion (*41*). As for *group_5289*, importing betaine has long been recognised as an important method of surviving in urinary tract infections (*42, 43*), a niche which *N. meningitidis* has long been known to have the ability to infect (*44*), and appears to be increasing in prevalence (*45*).

The GWES results have this far been discussed at gene level. Even though SpydrPick outputs links between specific sites, we recommend that the initial examination of the discovered links is kept at gene resolution, since fine-mapping the exact location of SNPs under selection in a GWES is typically very difficult. However, once a link between an interesting gene pair has been identified, one might still want to zoom in and look for further evidence of co-selection at SNP resolution. In particular, when an identified SNP is located in a protein-coding region, one might want to check if the SNP is synonymous or non-synonymous. As an illustrative example, we looked closer at the SNPs involved in the link between *trpE* and *group_5289*. While the SNP in the *group_5289* was found to be non-synonymous, resulting in an arginine to lysine mutation, the SNP in *trpE* was found to be synonymous at the protein-coding level. As synonymous mutations are not typically expected to be under selection, we scanned the surrounding region of the *trpE* site to look for a biologically more likely source of the signal. More specifically, using the SpydrPick output, we extracted all *trpE* sites that were in strong LD (measured by MI) with the original *trpE* site. Using the MI of the original link between *trpE* and *group_5289* as a threshold, we found 14 candidate SNPs located 36 - 676 bp from the original *trpE* site. Among these, we found one non-synonymous SNP, coding an aspartic acid to alanine mutation. Finally, to predict the functional effect of the amino acid substitutions, we used SNAP2 which outputs a value between −100 (completely neutral) and 100 (high functional effect) (*46*). The predicted effects of the *group_5289* and *trpE* mutations were 45 and 32, respectively, making both likely candidates for mutations under selection.

### Runtime

Calculating the MI values and running the ARACNE post-processing step for the *S. pneumoniae* alignment (with 3,042 sequences and 94,880 sites) took 2 hours using 8 threads on a laptop with Intel Core i7-6820HQ CPU. In comparison, it took over a week for SuperDCA to run direct coupling analysis on the same alignment using a single 20-core dual-socket compute node (*10*).

## DISCUSSION

The rapidly increasing availability of population-wide genome sequence data has boosted the potential for data-driven exploration of genetic variation associated with bacterial evolution. As a result, high-dimensional exploratory data analysis methods have become valuable tools for generating detailed hypotheses and identifying important targets for subsequent experimental work. For eukaryotes, genome-wide association studies (GWAS) have been the primary tool for this purpose for more than a decade, and more recent works have demonstrated the applicability and potential of GWAS also for bacteria (*29, 47, 48*). In addition to GWAS, the phenotype-free approach of genome-wide epistasis and co-selection studies (GWES) has recently emerged, and successfully been used to uncover mechanisms behind complex bacterial traits associated with survival, proliferation and virulence (*8–10*).

The main advantage of GWES lies in its unsupervised approach. It does not require the definition and measurement of a phenotype, yet it can reveal co-evolutionary patterns behind many different traits shaped by selection. Bacterial genomes of a single species are likely sampled from diverse micro-niches, which create unique selective pressures that vary over space and time. These can include immune pressures, nutrient availability, antibiotic use, or interactions within ecological communities. Links identified by GWES may represent multilocus adaptation to these micro-niches, which will create combinations of mutations that are maintained by selection. This adaptive process may be facilitated by epistatic interactions between loci but may also be driven by independent selection on sets of mutations that are additively beneficial in a particular niche. Co-evolutionary signals may also be maintained in a population if negative frequency dependent selection (NFDS) acts on the same traits. In fact, it has recently been suggested that NFDS acts to prevent antibiotic resistance genes sweeping to fixation in *S. pneumoniae* (bioRxiv: https://doi.org/10.1101/233957).

In this work, we introduced the model-free GWES method SpydrPick, which is parallelizable and scalable to pan-genome-wide alignments of many bacteria. To illustrate the output of a GWES, we introduced a modified version of the Manhattan plot, which has served as the main illustrative tool for exploring the output of GWAS. Experiments on both synthetic and real bacterial population sequence data demonstrated the accuracy and potential of our method. In particular, a genome-wide analysis of a mapping-based alignment of *S. pneumoniae* isolates showed that SpydrPick was able to accurately pick out previously discovered and validated signals of co-selection, as well as a novel link with a plausible biological explanation. In addition, a pan-genome-wide analysis of a Roary generated alignment of *N. meningitidis* isolates illustrated the potential of our method in an even more challenging data set, by identifying several interesting signals likely to originate from genes under selection. Similar to previous GWES methods, SpydrPick operates on SNP resolution trying to fine-map the co-selection signal to individual sites using only the co-variation pattern observed in the data. For any method, this task is very challenging and limited by several factors, including population structure, extent of LD and amount of available data. As illustrated by the identified *trpE* site in *N. meningitidis*, it is likely to be informative to check the surrounding region of the statistically linked sites to find the biologically most plausible source of the signal.

SpydrPick is conceptually very different to model-based DCA methods, which aim to fit a joint model over all SNPs, in that the pairwise interaction between two sites is evaluated independently of all other sites. This is similar in spirit to the approach by Cui et al. (*8*), who used Fisher’s exact test to scan for epistatic interactions among bi-allelic SNPs in a sample of *Vibrio parahaemolyticus* isolates. In contrast to our method, however, Cui et al. did not attempt to disentangle the direct interaction from the indirect interactions. In a recent hybrid approach, Gao et al. proposed filtering the data based on pairwise correlations and then fitting a joint model over the remaining sites in (*49*). The obvious advantage of a strict pairwise method, such as SpydrPick, is that its computational simplicity allows for scaling up to data sets beyond what is currently achievable by current DCA-based methods. In addition, and more importantly, recent numerical experiments on synthetic network models suggest that pairwise methods may be more accurate than the current state-of-the-art DCA-based methods in the high-dimensional setting (arXiv:1901.04345).

To distinguish between LD-mediated and non-LD-mediated links, we used a distance-based threshold with a rather conservative default value set to 10 kbp. As the background distribution will depend on multiple factors, such as type of organism, mode of recombination, population structure of the sample etc, it might be necessary to adjust the threshold value accordingly. This may involve running the analysis twice, where the output of the initial run is solely used to re-adjust the LD threshold parameter according to the drop in LD observed in the Manhattan plot, for example, see Fig 4b. A topic for future research will be to look into alternative and more sophisticated means for distinguishing between LD-mediated and non-LD-mediated links. This will be particularly important for alignments where a distance-based threshold cannot be used easily, for example in the analysis of the pan-genome. However, it might also open up opportunities for identifying signals of co-selection between closely located SNPs.

Another important topic for future research is to compare different techniques for adjusting for the population structure. In the work by Cui et al. (*8*), a subsample of 51 unrelated isolates was selected for the co-variation analysis. This corresponds to a hard reweighting technique where each weight is set to either zero or one, meaning that a collection of closely related isolates is represented by a single isolate. In contrast, the conceptual idea behind the soft reweighting technique used here can be thought of as taking the average over the same collection of isolates. The optimal technique for adjusting for the population structure will likely depend on certain properties in the data, for example, the level of clonality among the isolates.

GWES is a relatively new data-driven approach for detecting co-evolutionary patterns shaped by selection, and it is currently gaining traction in bacterial genomics due its wide applicability. GWES is by design phenotype-free, however, if one has access to relevant phenotype data, the output of a GWES can also be used to effectively reduce the number of tests in a follow-up epistatic GWAS (*50*). Given its accuracy and computational scalability, SpydrPick pushes the boundaries of existing GWES methods and promises to uncover a wealth of previously-undiscovered evolutionary signals in bacterial genomic data.

## AVAILABILITY

A multiple sequence alignment of the *S. pneumoniae* strains is available from the Dryad Digital Repository: https://datadryad.org/resource/doi:10.5061/dryad.gd14g. The *N. meningitidis* strains are available from the European Nucleotide Archive (ENA) and their accession numbers are given in Table S4. The SpydrPick software is available from the GitHub repository: https://github.com/santeripuranen/SpydrPick.

## SUPPLEMENTARY DATA

Supplementary Data are available at NAR online.

## FUNDING

This work was supported by the COIN Center of Excellence, Academy of Finland [grant number 251170 to JP, SP, MP and YX]; a Wellcome Trust PhD scholarship [grant number 204016 to GTH]; Wellcome Trust [grant number 098051 to L.S.B.]; a Sir Henry Wellcome Postdoctoral Fellowship [grant number 107378/Z/15/Z to CC]; European Research Council [grant number 742158 to J.C.].

## CONFLICT OF INTEREST

The authors declare that there are no conflicts of interests.

