## Supplemental Tables S1-S3 for "Genome-wide epistasis and co-selection study using mutual information"

| Parameter name | Parameter value |
|---|---|
| **SEED** | 1337, 2337, …, 10337 |
| **GENR** | 20000 |
| **LOLE** | 200000 |
| **NLOC** | 1 |
| **NBAC** | 1000 |
| **NPOP** | 20 |
| **MIGI** | 0 |
| **MIGR** | 0.005 |
| **MIGP** | 0.001 |
| **MUTR** | 2e-7 |
| **RECR** | 4 |
| **RECL** | 2000 |
| **RECA** | 18 |
| **RSEL** | 1 |
| **RORD** | 1 |
| **SUMI** | 20000 |
| **SEQS** | 0.05 |
| **SEQI** | 20000 |

**Table S1.** Parameter settings used for generating data with the Bacmeta simulator. The simulator parameters are described in detail in the Bacmeta input/output manual, which is available at Bitbucket: https://bitbucket.org/aleksisipola/bacmeta.

| Index | Position 1 | Gene information | Position 2 | Gene information | Distance | MI | MI* |
|---|---|---|---|---|---|---|---|
| 1 | 1,601,843 | SPN23F16620 (divIVA) – putative cell-division protein DivIVA | 1,891,918 | SPN23F19490 – putative uncharacterized protein | 290,075 | 0.472 | 0.445 |
| 2 | 292,881 | SPN23F03080 (pbp2x) – penicillin-binding protein 2x | 1,613,136 | SPN23F16740 (pbp2b) – penicillin-binding protein 2b | 901,060 | 0.469 | 0.405 |
| 3 | 292,944 | SPN23F03080 (pbp2x) – penicillin-binding protein 2x | 1,613,136 | SPN23F16740 (pbp2b) – penicillin-binding protein 2b | 901,123 | 0.448 | 0.426 |
| 4 | 1,601,803 | Intergenic (15 bp from divIVA) | 1,891,780 | SPN23F19490 – putative uncharacterized protein | 289,977 | 0.442 | 0.410 |
| 5 | 123,650 | Intergenic (32 bp from pspA) | 1,893,422 | Intergenic (upstream of ply) | 451,543 | 0.438 | 0.041 |
| 6 | 123,727 | SPN23F01290 (pspA) - pneumococcal surface protein PspA | 1,893,470 | Intergenic (upstream of ply) | 451,572 | 0.436 | 0.306 |
| 7 | 123,603 | Intergenic (79 bp from pspA) | 1,891,918 | SPN23F19490 – putative uncharacterized protein | 453,000 | 0.422 | 0.406 |
| 8 | 293,220 | SPN23F03080 (pbp2x) – penicillin-binding protein 2x | 1,613,164 | SPN23F16740 (pbp2b) – penicillin-binding protein 2b | 901,371 | 0.393 | 0.365 |
| 9 | 292,932 | SPN23F03080 (pbp2x) – penicillin-binding protein 2x | 331,929 | SPN23F03410 (pbp1a) – penicillin-binding protein 1a | 38,997 | 0.383 | 0.344 |
| 10 | 578,828 | SPN23F05840 – putative uncharacterized protein | 1,821,010 | SPN23F18740 – nicotinamide mononucleotide transporter | 979,133 | 0.352 | 0.006 |
| 11 | 292,959 | SPN23F03080 (pbp2x) – penicillin-binding protein 2x | 333,156 | SPN23F03410 (pbp1a) – penicillin-binding protein 1a | 40,197 | 0.351 | 0.341 |
| 12 | 292,821 | SPN23F03080 (pbp2x) – penicillin-binding protein 2x | 332,814 | SPN23F03410 (pbp1a) – penicillin-binding protein 1a | 39,993 | 0.341 | 0.291 |
| 13 | 1,601,874 | SPN23F16620 (divIVA) – putative cell-division protein DivIVA | 1,893,593 | Intergenic (upstream of ply) | 291,719 | 0.333 | 0.332 |
| 14 | 302,316 | SPN23F03150 (dexB) – glucan 1,6-alpha-glucosidase | 1,891,780 | SPN23F19490 – putative uncharacterized protein | 631,851 | 0.324 | 0.285 |
| 15 | 331,983 | SPN23F03410 (pbp1a) – penicillin-binding protein 1a | 1,613,323 | SPN23F16740 (pbp2b) – penicillin-binding protein 2b | 939,975 | 0.297 | 0.235 |
| 16 | 294,490 | SPN23F03090 (mraY) – phospho-N-acetylmuramoyl-pentapeptide-transferase | 1,613,323 | SPN23F16740 (pbp2b) – penicillin-binding protein 2b | 902,482 | 0.285 | 0.241 |

**Table S2.** Top 16 SpydrPick hits in *S. pneumoniae* population. The pairs are ordered by their mutual information (MI). MI* refers to mutual information calculated from sequences for which neither position in the pair contains a gap.

| Index | Position 1 | Gene information | Position 2 | Gene information | Mock dist. | MC58 dist. | FAM18 dist. | MI | MI* |
|---|---|---|---|---|---|---|---|---|---|
| 1 | 8,651,268 | group_8012 – hyp./signal peptide/WD40 | 9,532,354 | group_9383 – hyp./phage ass./transcript. reg. | 881,086 | 642 | - | 1.038 | 0.656 |
| 2 | 8,651,268 | group_8012 – hyp./signal peptide/WD40 | 9,532,233 | group_9382 – DNA bind./transcript. reg./phage repr. | 880,965 | 373 | - | 1.037 | 0.655 |
| 3 | 9,532,354 | group_9383 – hyp./phage ass./transcr. regul. | 9,532,636 | group_9384 – DNA binding/phage resp. | 282 | 226 | - | 1.028 | 0.639 |
| 4 | 8,518,065 | group_7779 – hyp./methyltransferase | 9,469,266 | group_9227 – hyp./recombinase family | 951,201 | 427 | 427 | 1.026 | 0.657 |
| 5 | 7,120,760 | group_5567 – d-lactate dehydr./phage ass./Myb/AlpA | 7,123,730 | group_5573 – hyp./lactate dehydrog./phage ass. | 2,970 | 207 | 207 | 1.011 | 0.665 |
| 6 | 8,922,542 | group_8344 – DUF2303 | 9,469,266 | group_9227 – hyp./recombinase family | 546,724 | 1,306 | 1,306 | 1.010 | 0.617 |
| 7 | 7,123,778 | group_5573 – hyp./lactate dehydrog./phage ass. | 8,517,646 | group_7778 – hyp./signal peptide | 1,393,868 | 919 | 920 | 1.007 | 0.655 |
| 8 | 8,360,604 | group_7554 – DNA binding/phage ass./BcepMu gp16 | 8,651,268 | group_8012 – hyp./signal peptide/WD40 | 290,664 | 1,106 | 783,179 | 1.002 | 0.640 |
| 9 | 8,360,539 | group_7554 – DNA binding/phage ass./BcepMu gp16 | 9,532,233 | group_9382 – DNA bind.transcript. reg./phage repr. | 1,171,694 | 733 | - | 1.001 | 0.638 |
| 10 | 6,169,347 | group_4419 – DUF3102 /phage ass./Gp10 | 8,651,268 | group_8012 – hyp./signal peptide/WD40 | 2,481,921 | 2,328 | 781,957 | 0.996 | 0.602 |
| 11 | 8,517,646 | group_7778 – hyp./signal peptide | 8,922,542 | group_8344 – DUF2303 | 404,896 | 285 | 287 | 0.972 | 0.560 |
| 12 | 8,517,695 | group_7778 – hyp./signal peptide | 9,469,162 | group_9227 – hyp./recombinase family | 951,467 | 1,591 | 1,593 | 0.946 | 0.539 |
| 13 | 4,213,372 | group_2190 – DUF4760/phage ass./membr./signal | 8,517,562 | group_7778 – hyp./signal peptide | 4,304,190 | 3,438 | 3,449 | 0.927 | 0.544 |
| 14 | 21,353 | p17_acnA – aconitate hydratase | 10,703,638 | prpF – 2-methylaconitate cis-trans isomerase | 10,682,285 | 3,409 | 1,522 | 0.925 | 0.621 |
| 15 | 8,265,404 | group_7339 – mRNA iterferase/pemK | 8,517,695 | group_7778 – hyp./signal peptide | 252,291 | 4,254 | 4,265 | 0.910 | 0.507 |
| 16 | 7,120,760 | group_5567 – d-lactate dehydr./phage ass./Myb/AlpA | 8,061,276 | group_6987 – hyp./integral membrane/phage ass. | 940,516 | 6,047 | 27,470 | 0.909 | 0.530 |
| 17 | 121,999 | besA – ferri-bacillibactin esterase | 478,005 | fpvA – ferripyoverdine receptor | 356,006 | 62,447 | 63,514 | 0.895 | 0.603 |
| 18 | 7,845,699 | group_6635 – protease HtpX hom./heat shock HtpX | 7,846,128 | group_6636 – orotidine 5'-phosphate decarboxylate | 429 | 2,227 | 2,223 | 0.886 | 0.570 |
| 19 | 122,450 | besA – ferri-bacillibactin esterase | 478,501 | fpvA – ferripyoverdine receptor | 356,051 | 62,447 | 63,514 | 0.884 | 0.593 |
| 20 | 6,913,189 | group_5289 – hyp./betaine transporter | 11,186,468 | trpE – anthranilate synthase component I | 4,273,279 | 361,849 | 722,196 | 0.882 | 0.511 |
| 21 | 122,230 | besA – ferri-bacillibactin esterase | 478,690 | fpvA – ferripyoverdine receptor | 356,460 | 62,447 | 63,514 | 0.881 | 0.587 |
| 22 | 122,185 | besA – ferri-bacillibactin esterase | 478,408 | fpvA – ferripyoverdine receptor | 356,223 | 62,447 | 63,514 | 0.879 | 0.583 |
| 23 | 6,120,411 | group_4367 – Hemerythrin cation binding | 6,462,382 | group_4800 – Diadenosine tetraphosphatase | 341,971 | 559 | 559 | 0.866 | 0.645 |
| 24 | 8,638,454 | group_7992 – hyp./lipoic acid synthase/DUF4124 | 10,341,339 | Mpl – murein peptide ligase | 1,702,885 | 367 | 367 | 0.865 | 0.622 |
| 25 | 8,061,354 | group_6987 – hyp./integral membrane/phage ass. | 9,614,083 | group_951 – hyp./ATP binding/peptidase | 1,552,729 | 644 | 23,875 | 0.863 | 0.652 |
| 26 | 8,061,354 | group_6987 – hyp./integral membrane/phage ass. | 9,469,162 | group_9227 – hyp./recombinase family | 1,407,808 | 3,330 | 24,750 | 0.861 | 0.461 |
| 27 | 7,020,647 | group_5442 – transposase/IS-200 | 11,038,305 | sucB – dihydrolipoamide succinyltransferase | 4,017,658 | 5,430 | 954,229 | 0.853 | 0.510 |
| 28 | 5,536,716 | group_3579 – peptidoglycan domain/lysozyme | 8,651,006 | group_8010 – hyp./lipoprotein/membrane | 3,114,290 | 2,042 | - | 0.841 | 0.657 |

**Table S3.** Top 28 SpydrPick hits in the *N. meningitidis* population. The pairs are ordered by their mutual information (MI). MI* refers to mutual information calculated from sequences for which neither position in the pair contains a gap.
